# Spatio-temporal dynamics of dengue in Brazil: seasonal travelling waves and determinants of regional synchrony

**DOI:** 10.1101/475335

**Authors:** Mikhail Churakov, Christian J. Villabona-Arenas, Moritz U.G. Kraemer, Henrik Salje, Simon Cauchemez

## Abstract

Dengue continues to be the most important vector-borne viral disease globally and in Brazil, where more than 1.4 million cases and over 500 deaths were reported in 2016. Mosquito control programmes and other interventions have not stopped the alarming trend of increasingly large epidemics in the past few years.

Here, we analyzed monthly dengue cases reported in Brazil between 2001 and 2016 to better characterize the key drivers of dengue epidemics. Spatio-temporal analysis revealed recurring travelling waves of disease occurrence. Using wavelet methods, we characterised the average seasonal pattern of dengue in Brazil, which starts in the western states of Acre and Rondônia, then travels eastward to the coast before reaching the northeast of the country. Only two states in the north of Brazil (Roraima and Amapá) did not follow the countrywide pattern and had inconsistent timing of dengue epidemics throughout the study period.

We also explored epidemic synchrony and timing of annual dengue cycles in Brazilian regions. Using gravity style models combined with climate factors, we showed that both human mobility and vector ecology contribute to spatial patterns of dengue occurrence.

This study offers a characterization of the spatial dynamics of dengue in Brazil and its drivers, which could inform intervention strategies against dengue and other arboviruses.

**Author summary:** In this paper we studied the synchronization of dengue epidemics in Brazilian regions. We found that a typical dengue season in Brazil can be described as a wave travelling from the western part of the country towards the east, with the exception of the two most northern equatorial states that experienced inconsistent seasonality of dengue epidemics.

We found that the spatial structure of dengue cases is driven by both climate and human mobility patterns. In particular, precipitation was the most important factor for the seasonality of dengue at finer spatial resolutions.

Our findings increase our understanding of large scale dengue patterns and could be used to enhance national control programs against dengue and other arboviruses.

## Introduction

Dengue fever is a mosquito-borne viral disease that causes serious health and economic burden in tropical and sub-tropical regions [1]. Population growth, urbanisation, international travel and changes in climate patterns have led to significant geographic expansion and continued rise in dengue cases [2,3]. In Brazil, dengue is an important public health concern with worsening societal and economic burden [4–7]. Large territories, various climate types, as well as heterogeneities in demographics, land use and urban development contribute to complex epidemic dynamics. In 2016, Brazilian public health authorities reported more than a million probable cases and over 600 deaths to the World Health Organization (WHO), while the number of apparent dengue infections was estimated at around 5 million per year [8]. With no effective dengue treatment, control of the disease is limited but mostly done through vector control interventions [9]. To design appropriate response strategies, a detailed understanding of the dynamics of dengue spread at the country level is necessary [10]. Applications of such knowledge might well expand to other arboviruses transmitted by the same vector [11–13].

Dengue has previously been shown to exhibit ‘travelling wave’ type dynamics [14,15], i.e. when regional epidemic timing has a distinct spatial structure. However, these studies have largely focused on Southeast Asian settings. It remains unclear, to which extent pan-national waves are also observed in South America, where dengue re-emergence occurred more recently [4]. On a more local scale, it has been shown that in

Brazil seasonal waves of dengue spread from metropolitan areas towards smaller cities in the same region [16]. In addition, phylodynamic approaches showed that there are repeated introductions of dengue viruses from northern to southern Brazilian states [17,18].

The reasons why communities may experience a lag in dengue epidemics compared to their neighbours can be attributed to multiple factors. Synchonry of regional epidemics can be mediated by climate drivers such as temperature and rainfall, as has previously been observed between different countries within Southeast Asia [15,19,20]. The movement of infectious people between communities might also provide wave-like dynamics across settings [21–23]. The contributions of climate and human mobility in describing the synchrony in dengue epidemics between locations have rarely been considered alongside each other.

Here, we analysed 16 years of data on reported dengue cases in Brazil to examine seasonal travelling waves across the country. We characterised the average seasonal pattern and, using gravity style models for human mobility combined with climate variables, identified key factors associated with the observed patterns of dengue occurrence.

## Data and methods

### Dengue cases data

Cases data were obtained from the Notifiable Diseases Information System (SINAN, Sistema de Informação de Agravos de Notificação) via the Health Information Department (DATASUS, Departamento de Informática do Sistema Único de Saúde) run by the Brazilian Ministry of Health. Monthly dengue data were extracted from January 2001 to August 2016 for each municipality (n=5570). Cases were confirmed by clinical and epidemiological evidence, and approximately 30% of them were also laboratory-confirmed [24].

### Population data

Municipality level human population data were retrieved from the Brazilian Institute of Geography and Statistics (IBGE, Instituto Brasileiro de Geografia e Estatística) for the year 2014 (http://www.ibge.gov.br/home/estatistica/populacao/estimativa2014/estimativa_dou.shtm).

### Climate data

Precipitation data were obtained from the Climate Prediction Center's (CPC) rainfall data for the world (1979 to present, 50 km resolution) via raincpc R package [25] and aggregated for each Brazilian municipality over the study period. We also retrieved data on mean monthly surface temperature from Reanalysis CFSR model [26] (reference: CREATE-IP.reanalysis.NOAA-NCEP.CFSR.atmos.mon, data node: esgf.nccs.nasa.gov).

### Vector suitability data

We obtained average monthly estimates of vector suitability for each Brazilian municipality using the modelling outputs from [27,28]. Mosquito occurrence data were fitted to annual data and covariates (*i.e.* time varying temperature-persistence suitability, relative humidity, precipitation and a static urban vs. rural covariate) and then subsequently re-applied to the same covariates at a monthly level. Maps were produced at a 5 x 5 kilometre resolution, aggregated to the municipality level and the mean value was used for our model. For consistency, we rescaled the monthly suitability values, so that the sum of all monthly maps equalled the annual mean map.

### Characterising spatial patterns of dengue: wavelet transform and phase angles

We used `biwavelet` package [29] in R with Morlet function as a wavelet base for the analysis of longitudinal dengue data. Original case series for each state were log-transformed and scaled to zero mean and unit variance. The outputs of the wavelet transform of each of the time series consisted of: the local wavelet power spectrum, which allows for inspection of time-frequency distribution and detection of predominant signal components for particular time periods; and the corresponding phase angles, which can be used to assess the speed of wave propagation for a particular period.

The annual signal dominated in state level case series (individual power spectra for each state can be found in S3–S5 Figs), and to explore it further we extracted phase angles for the annual component. The next step was to obtain pairwise phase angle differences between states that indicate for each point in time whether a state is ahead or behind another one in terms of recurrent annual waves. Consequently, for each state we obtained the average phase difference from the other 26 states. The mean value of this phase difference over time was used to produce a map of annual phase lags between states. Phase differences were interpreted as time units as for the regular annual signal phase angle changes from -π to +π in 1 year, which makes the phase lag of 1 radian equivalent to approximately 2 months. Hence, the map of annual phase lags represents the average ordering of states in terms of dengue wave arrival times.

### Epidemic synchrony and annual phase coherence

We considered correlations between regional time series by computing the Pearson correlation coefficient of raw case series (epidemic synchrony) and annual phase angles (annual phase coherence) for each pair of regions. Then, these measures were summarised by the nonparametric spline covariance function (implemented in ncf R package [30]) to assess how they depend on the distance. Normally, they are higher between neighbouring regions that experience synchronized dengue epidemics and, therefore, could serve as useful descriptors of travelling waves. Phase coherence measures the relative timing of seasonal epidemics while epidemic synchrony indicates how their relative amplitudes covary [31]. In other words, annual phase coherence describes lags between annual signals (i.e. seasonality only), while epidemic synchrony also accounts for other frequency components as well as their amplitudes (other synchronised events with various periodicity).

### Identifying determinants of dengue synchrony

We developed a set of statistical models to characterize the determinants of epidemic synchrony and annual phase coherence [22]. The models consider covariates such as demography, human mobility, climate factors (surface temperature and precipitation) and disease vector suitability. They are defined by the following equation:

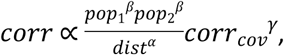

where *corr* is the correlation measure (i.e. either epidemic synchrony or phase coherence), *pop1* and *pop2* are the population sizes of the two regions, *dist* is the distance between the two regions, *corr_cov_* is the correlation of climate or environmental covariance time series (temperature, precipitation or vector suitability) between the two regions. The first part of the equation 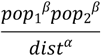 is a standard gravity model that captures the contribution of human mobility and demography [32,33] while the second part *corr_cov_*^γ^ captures the fact that similar climate profiles may also partly explain synchrony in epidemic time series.

Different model variants were considered by fixing various combinations of parameters *α*, *β* and *γ* to zero. Starting with the null model (*α = 0*, *β = 0, γ = 0*), distance only (*β = 0, γ = 0*) and population only (*α = 0, γ = 0*) models, original gravity model (*γ = 0*), and for each climate factor: model without human mobility (*α = 0*, *β = 0*) and the full model.

The exponents *α*, *β* and *γ* were estimated using a linear regression of the log-transformed form of the original equation:

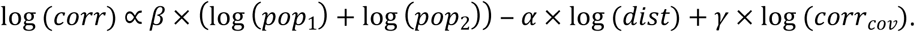

We excluded data on region pairs for which we had negative correlations to allow for log-transformation. We used R^2^ and Akaike’s Information Criterion (AIC) for model comparison, and also generated bootstrapped confidence intervals for the parameters by resampling the location pairs with replacement 500 times.

### Spatial resolution

We used different spatial data aggregation levels given that a large number of municipalities had very few cases for certain seasons. We used three official administrative levels: state level (n = 27, 26 states and the federal district Brasilia, hereinafter referred to as state), mesoregion level (n = 137) and microregion level (n = 558). In addition, we performed our analysis using alternative Urban-Regional divisions (https://ww2.ibge.gov.br/home/geociencias/geografia/default_divisao_urbano_regional.shtm): Urban-1 (n = 14), Urban-2 (n = 161) and Urban-3 (n = 482).

In this paper, we present results of the wavelet analysis on the state level, primarily for better visual representation. Analyses of epidemic synchrony and annual phase coherence are presented on the Urban-2 level (n = 161), to allow for sufficient variation in distance between pairs of regions. The results on all the six available levels can be found in the Supplementary Material.

## Results

### Annual patterns of dengue outbreaks

Over the study period from January 2001 to August 2016, there were a total of over 8.5 million reported dengue cases, with an average of 500,000 cases per year. During the study period, the highest risk of dengue (i.e. the highest number of total reported cases per capita) was for Acre, Mato Grosso do Sul and Goiás (Fig 1A). These states had an average of 101, 69 and 66 annual reported cases per 10,000 inhabitants, respectively. The southern states Santa Catarina and Rio Grande do Sul had very few dengue cases compared to other states, with an annual average below one case per 10,000 inhabitants. We observed a strong seasonal pattern with the majority of cases occurring between December and June (Fig 1B, 1C) and that dengue typically peaks slightly earlier in the western states compared to the eastern ones (Fig 1B, 1D).

**Fig 1.**
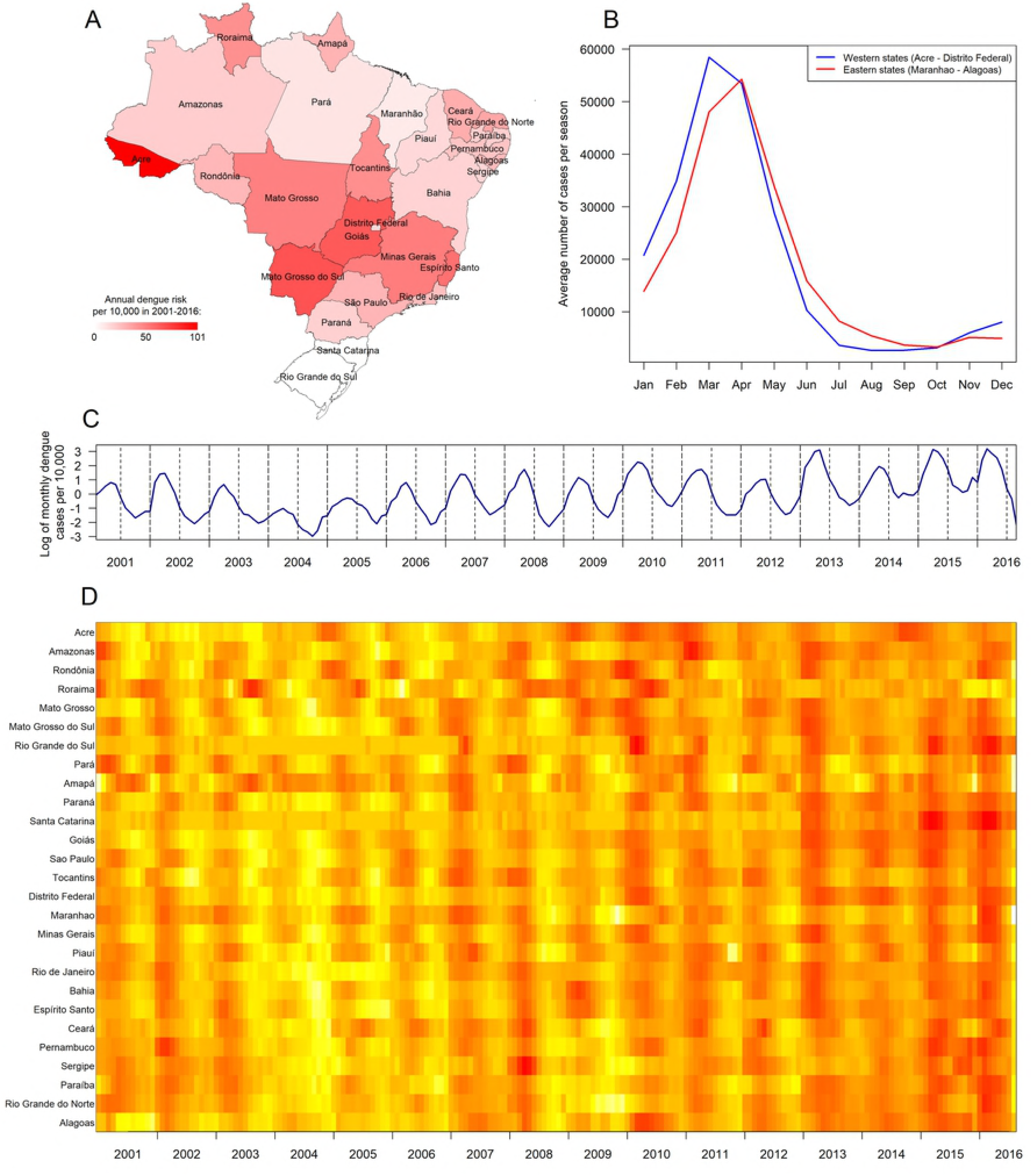
Summary of the spatio-temporal dynamics of dengue in Brazil. (A) Average annual dengue risk per 10,000 inhabitants in 2001–2016 for each state. Administrative boundaries were obtained from GADM (https://gadm.org). (B) Average number of cases for each month in the eastern and the western states, relatively to Distrito Federal. (C) Log of monthly dengue cases per 10,000 inhabitants in Brazil. (D) Heat map of log-normalised case series for Brazilian states sorted by longitude: west (top) to east (bottom). Colours were scaled for each state independently so that yellow indicates the lowest number of dengue cases and red indicates the maximum.

### Average seasonal spatial pattern

We detected a significantly predominant annual component of dengue time series throughout the whole study period for most of the states (S1 Fig) apart from Roraima and Amapá (two northern states that are also the least populated in Brazil) that had additional strong components with a period of 1.5–2 years, and the southern states of Rio Grande do Sul and Santa Catarina that had very few dengue cases for several years.

Figure 2A shows, for each state, the median and 95% range of annual phase lags from other states, which represent delays of individual state dengue waves from the nationwide annual waves. Average phase lags enabled us to identify the average seasonal pattern of dengue in Brazil (Fig 2B), which starts in the western states, travels to the highly densely populated southeastern states of São Paulo and Rio de Janeiro, and then reaches the northeast of the country. We found that Roraima and Amapá, two states in the north, had inconsistent annual phase lags over time (S2 Fig) suggesting that they had poor synchrony with the rest of the country and did not follow the aforementioned wave pattern. We observed similar spatial patterns when we aggregated data at different spatial resolutions (S3 Fig).

**Fig 2.**
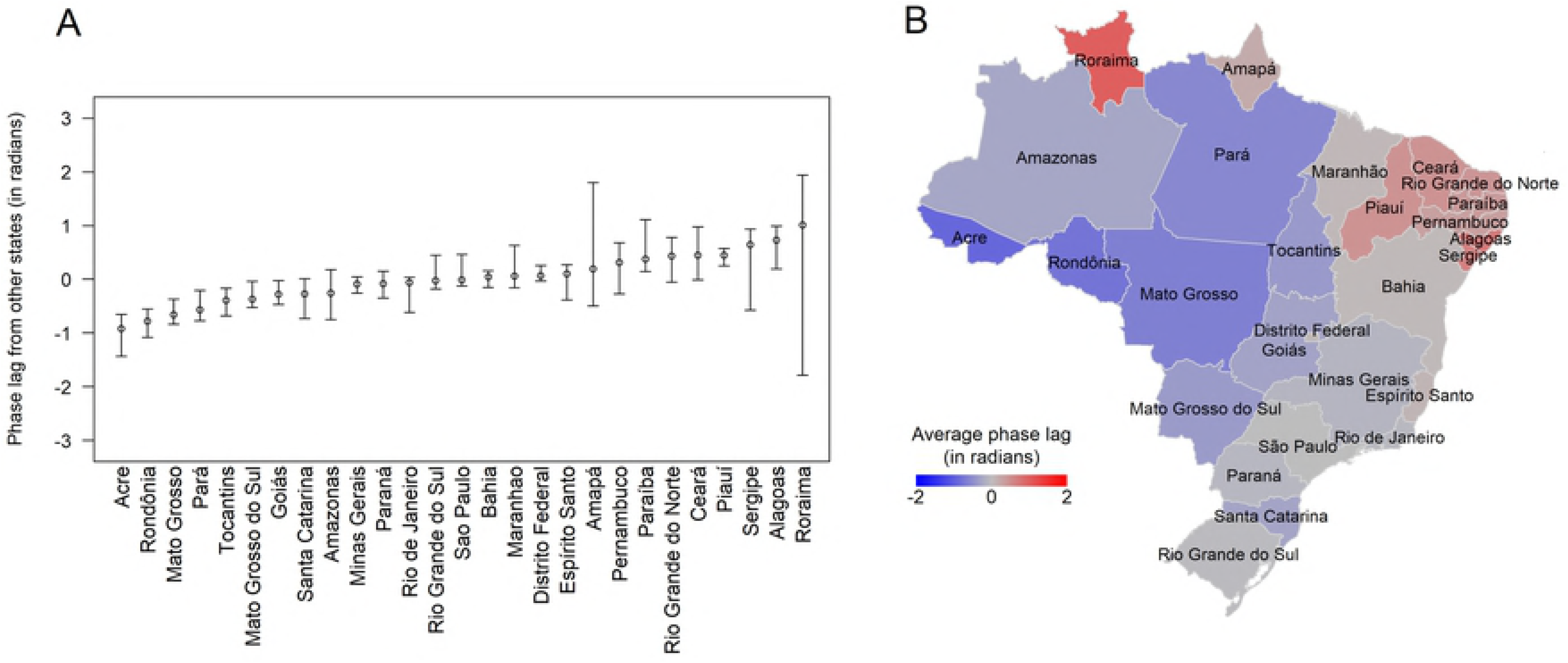
Average phase lags of seasonal dengue between Brazilian states. (A) For each state, median and 95% range of annual phase lags from other states over the study period. States were ordered by their median phase lag. (B) Map of the relative timing of annual dengue waves, which was defined using the average annual phase lag of each state from every other state. Administrative boundaries were obtained from GADM (https://gadm.org).

### Epidemic synchrony and annual phase coherence

We explored correlations between dengue time series in different regions. Both epidemic synchrony and phase coherence were higher for closer regions and declined with distance (Fig 3). For the Urban-2 (n = 161) spatial level, epidemic synchrony reached the average countrywide correlation at approximately 1,260 kilometres (Fig 3A). This synchrony length represents a substantial part of Brazil’s dimensions as the country extends 4,395 kilometres north to south and 4,319 kilometres west to east. The coherence length had a higher value of 1,590 kilometres (Fig 3B), suggesting that agreement in dengue seasonality spreads further than correlations of epidemic curves.

**Fig 3.**
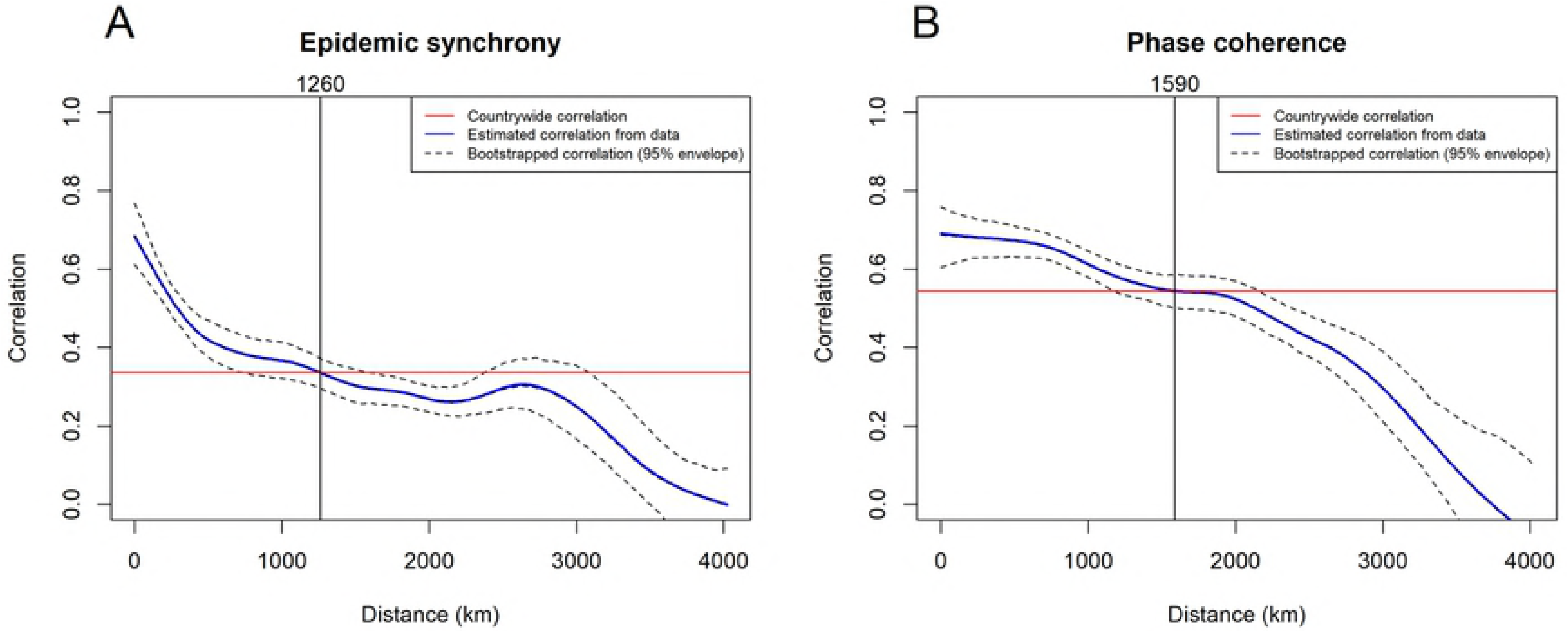
Epidemic synchrony and annual phase coherence between Brazilian Urban-2 regions. Epidemic synchrony (A) and annual phase coherence (B) summarised using nonparametric spline covariance function. Solid blue line describes the mean pairwise correlation from the data and the dotted lines represent the 95% envelope for bootstrapped correlations of case and annual phase angle time series, respectively. Red line indicates global countrywide correlation.

We also looked at epidemic synchrony and annual phase coherence at other spatial levels (S4 and S5 Figs, respectively) and found that both synchrony and coherence lengths tend to decrease for smaller spatial resolutions and stabilise at 1,240 km and 1,500 km.

### Determinants of dengue epidemic synchrony and annual phase coherence

We built a suite of models investigating potential determinants of dengue epidemic synchrony (Table 1). The classical gravity model (model 4) that accounted only for distance and the product of population sizes of regions captured a part of variation in epidemic synchrony (9.6%) which was higher than those of vector suitability (1.5%, model 6), precipitation (no variance explained, model 8) and average temperature (3.8%, model 10). The fit was further improved by combining the gravity model with correlations of vector suitability (model 5), precipitation (model 7) and average temperature (model 9), raising the explained variance up to 11%, 14% and 11%, respectively. We found that estimates of human mobility parameters were consistent between different models, regardless of additional variables.

**Table 1.** Gravity style model fitting for epidemic synchrony between Urban-2 regions.

| | Model | $K$<br>mean (CI) | $\alpha$ | $\beta$ | $\gamma$ | $R^2$ | AIC |
| --- | --- | --- | --- | --- | --- | --- | --- |
| 1 | $K$ | 0.26 (0.26-0.27) | - | - | - | 0 | 32458 |
| 2 | $\frac{K}{dist^\alpha}$ | 2.9 (2.5-3.3) | 0.71 (0.70-0.73) | - | - | 0.079 | 31433 |
| 3 | $K(pop_i pop_j)^\beta$ | 0.028 (0.021-0.037) | - | 1.09 (1.08-1.10) | - | 0.018 | 32229 |
| 4 | $K \frac{(pop_i pop_j)^\beta}{dist^\alpha}$ | 0.34 (0.24-0.46) | 0.72 (0.70-0.73) | 1.08 (1.07-1.09) | - | 0.096 | 31212 |
| 5 | $K \frac{(pop_i pop_j)^\beta}{dist^\alpha} corr_{suit}^\gamma$ | 0.38 (0.27-0.52) | 0.71 (0.69-0.72) | 1.08 (1.07-1.09) | 1.00 (0.97-1.03) | 0.11 | 28906 |
| 6 | $K corr_{suit}^\gamma$ | 0.28 (0.27-0.28) | - | - | 1.13 (1.10-1.16) | 0.015 | 30127 |
| 7 | $K \frac{(pop_i pop_j)^\beta}{dist^\alpha} corr_{precip}^\gamma$ | 0.73 (0.51-1) | 0.62 (0.61-0.64) | 1.08 (1.07-1.10) | 0.92 (0.90-0.93) | 0.14 | 23615 |
| 8 | $K corr_{precip}^\gamma$ | 0.28 (0.28-0.29) | - | - | 1.07 (1.06-1.09) | -0.01 | 25150 |
| 9 | $K \frac{(pop_i pop_j)^\beta}{dist^\alpha} corr_{temp}^\gamma$ | 0.21 (0.15-0.28) | 0.76 (0.74-0.77) | 1.09 (1.08-1.10) | 1.11 (1.08-1.13) | 0.11 | 29425 |
| 10 | $Kcorr_{temp}^{\gamma}$ | 0.31 (0.3-0.32) | - | - | 1.25 (1.22-1.28) | 0.038 | 30276 |

The same analysis for annual phase coherence (see Table 2) revealed that while the gravity model performed even better (14% of variance explained), incorporation of other factors was still beneficial. The best fit (28% of variance explained) was for model 8 that accounted for human mobility and precipitation.

**Table 2.**
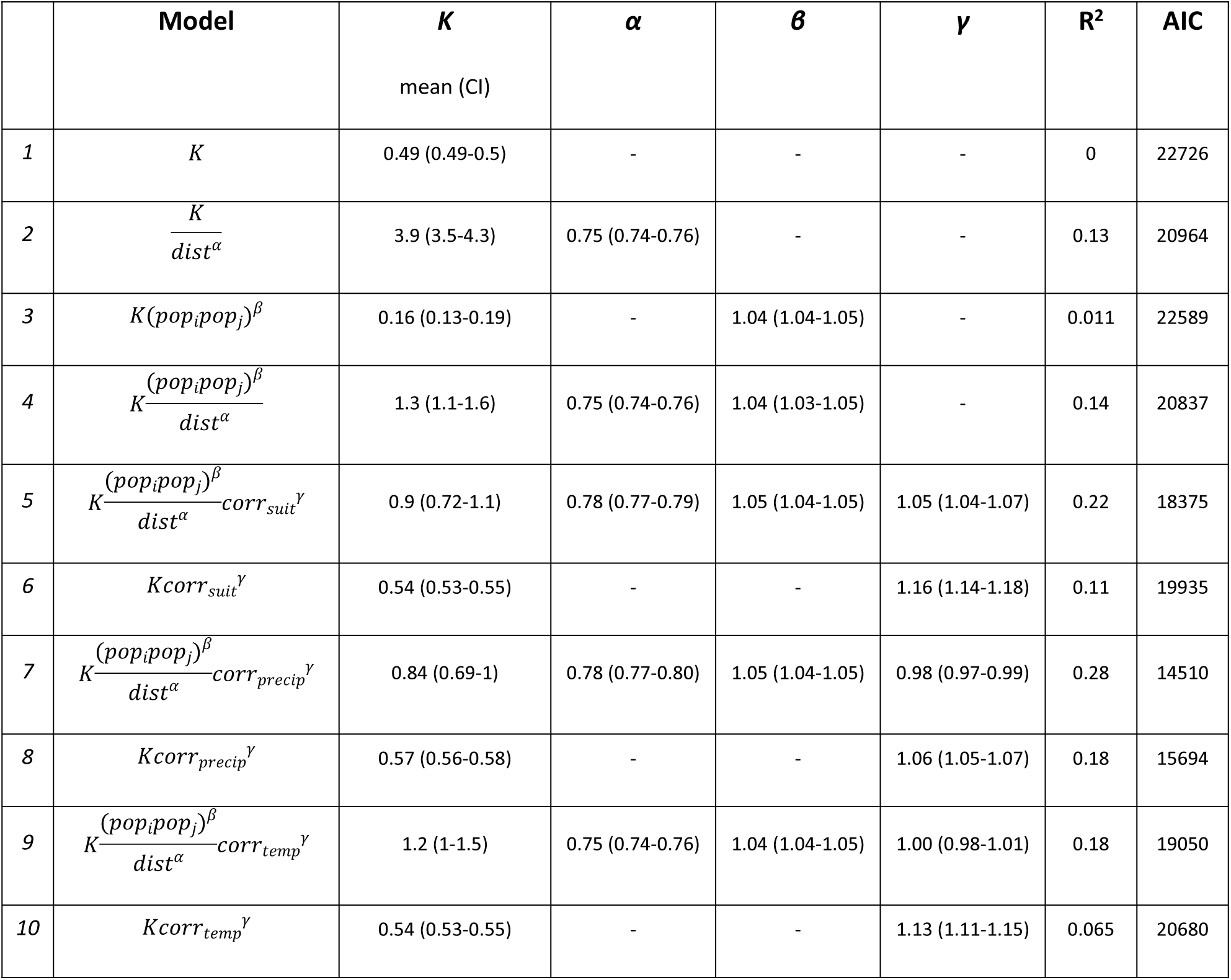
Gravity style model fitting for annual phase angle correlation between Urban-2 regions.

We explored performance of the statistical models for other spatial levels (see Fig 4) and found that overall model 8 (gravity model combined with precipitation) was the best in terms of variance explained. The classical gravity model (compared to climate factors) explained the majority of variance in epidemic synchrony across most of spatial scales (Fig 4A). However, at smaller scales, precipitation contributed the most for coherent timing of annual epidemics (Fig 4B).

**Fig 4.**
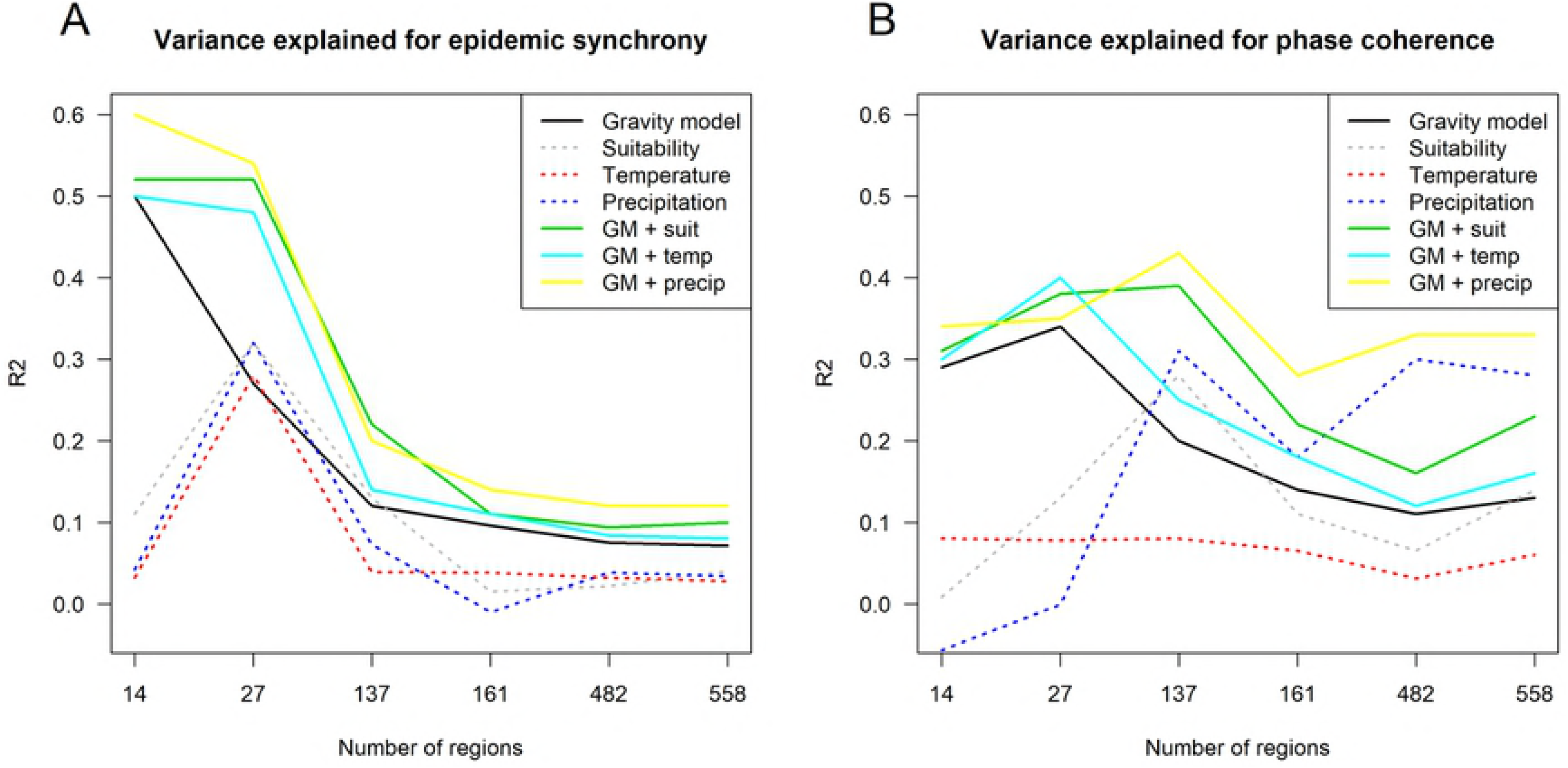
Variance explained for models of epidemic synchrony and annual phase coherence. R^2^ for epidemic synchrony (A) and annual phase coherence (B) predicted by models 4–10 depending on the spatial scale considered.

## Discussion

Here, we analysed longitudinal time series of dengue cases reported in Brazil. We found a consistent seasonal travelling wave type pattern of dengue occurrence and underscored human mobility in combination with climate variables as the potential determinants of spread.

Our findings revealed the presence of seasonal travelling wave that starts in the western states, then travels to the east and reaches the northeast at the end of a typical dengue season. Overall the timing of epidemic peaks between states was largely consistent between seasons. This provides a level of predictability for public health planning: particularly, in terms of preparedness of health care facilities to allocate enough resources for timely treatment of severe dengue cases. In particular, the epidemics in the more populous states along the coast appeared to peak relatively late compared to other states, potentially providing an opportunity for control programs to optimise the timing and placement of interventions.

The epidemiology of dengue in the two most northern states of the country (Roraima and Amapá) appeared different from the rest of the country, as they did not follow the overall seasonal travelling wave pattern. This could be explained by the equatorial climate with favourable conditions for dengue transmission throughout the year, while epidemics in other regions are highly restricted by seasonal variation in temperature and precipitation.

We characterized the spatial correlation between dengue cases series of each pair of regions depending on distance and estimated the extent of synchronised epidemics in terms of timing (annual phase coherence) and intensity (epidemic synchrony). Our results are in line with previously published ones that reported the spatial correlation of dengue cases between pairs of municipalities to decay with distance [34]. Our findings are also similar to a travelling wave of influenza identified in Brazil: from equatorial regions with low population to highly populous temperate regions [35].

Gravity models that used distance and population size as markers of human mobility explained a major part of the correlation in dengue epidemics. However, we found that incorporating information on the correlation in climate markers of vector ecology substantially improved the fit. Future research on relative contributions of vector ecology, human demographics, population mobility and host immunity could further increase our understanding of spatial dynamics of dengue and similar arboviruses.

Our findings suggest that precipitation was more important than human mobility for the seasonality of dengue at the mesoregion and finer spatial levels. Contrastingly, across scales human mobility better explained season-specific and region-specific features attributed to non-annual components and epidemic size effects. Contribution of human mobility approximated via gravity model for both epidemic synchrony and phase coherence was declining with finer spatial resolution. This might be suggesting that gravity models are failing to capture fine-scale human mobility and models that use empirical data are likely to perform better.

This study has several limitations. Firstly, we described the averaged seasonal pattern but did not consider season-specific features or anomalies (e.g. due to El Niño–Southern Oscillation). Incorporating these elements may further improve model performance. Secondly, in wavelet analysis we ignored information on amplitudes and considered only annual components of dengue case series. This was done considering the timing of seasonal dengue rather than its epidemic size and severity. Lastly, dengue surveillance system in Brazil is not entirely accurate for capturing all dengue cases, there might be regional biases, underestimation and overestimation issues [36,37]. However, as we mainly focused on the timing of dengue waves and on correlations of case series, these potential flaws in the cases data should not affect the presented results.

Our findings have potential implications for other circulating arboviruses in Brazil. Since they share the same vector, they could spread in similar ways, although differences between the viral characteristics (e.g. transmissibility, virulence, etc.) should be taken into account, as well as the influence of population immunity.

## Supporting information

**S1 Fig. Wavelet spectra for each Brazilian state.** (PDF)

**S2 Fig. Monthly phase lags (from other states) of each Brazilian state**. Roraima and Amapá were the least consistent while other states had similar phase lags throughout the study period. (PDF)

**S3 Fig. Spatial structure of average phase lags of seasonal dengue in Brazilian regions.** (A) state level, (B,C) meso- and micro-regions, (D,E,F) urban subdivisions. Administrative boundaries for Brazilian municipalities were obtained from IBGE (https://www.ibge.gov.br/geociencias-novoportal/cartas-e-mapas.html), along with shapefiles for Urban-Regional divisions (https://ww2.ibge.gov.br/home/geociencias/geografia/default_divisao_urbano_regional.shtm). (PNG)

**S4 Fig. Epidemic synchrony of seasonal dengue in Brazilian regions.** (A) state level, (B,C) meso- and micro-regions, (D,E,F) urban subdivisions. (PNG)

**S5 Fig. Phase coherence of seasonal dengue in Brazilian regions**. (A) state level, (B,C) meso- and micro-regions, (D,E,F) urban subdivisions. (PNG)

